# A comprehensive motifs-based interactome of the C/EBPα transcription factor

**DOI:** 10.1101/2020.12.28.424569

**Authors:** Evelyn Ramberger, Valeria Sapozhnikova, Elisabeth Kowenz-Leutz, Karin Zimmermann, Nathalie Nicot, Petr V. Nazarov, Daniel Perez-Hernandez, Ulf Reimer, Philipp Mertins, Gunnar Dittmar, Achim Leutz

**Affiliations:** Max Delbrück Center for Molecular Medicine in the Helmholtz Association, Robert-Rössle-Strasse 10, 13125 Berlin, Germany; Quantitative Biology Unit, Luxembourg Institute of Health, Luxembourg; JPT Peptide Technologies GmbH, Volmerstrasse 5, 12489 Berlin, Germany; Institute of Biology, Humboldt University of Berlin, 10115 Berlin, Germany

**Keywords:** intrinsic disorder, CEBPA interactome, BioID, peptide array, mass-Spectrometry

## Abstract

The pioneering transcription factor C/EBPα coordinates cell fate and cell differentiation. C/EBPα represents an intrinsically disordered protein with multiple short linear motifs and extensive post-translational side chain modifications (PTM), reflecting its modularity and functional plasticity. Here, we combined arrayed peptide matrix screening (PRISMA) with biotin ligase proximity labeling proteomics (BioID) to generate a linear, isoform specific and PTM-dependent protein interaction map of C/EBPα in myeloid cells. The C/EBPα interactome comprises promiscuous and PTM-regulated interactions with protein machineries involved in gene expression, epigenetics, genome organization, DNA replication, RNA processing, and nuclear transport as the basis of functional C/EBPα plasticity. Protein interaction hotspots were identified that coincide with homologous conserved regions of the C/EBP family and revealed interaction motifs that score as molecular recognition features (MoRF). PTMs alter the interaction spectrum of multi-valent C/EBP-motifs to configure a multimodal transcription factor hub that allows interaction with multiple co-regulatory components, including BAF/SWI-SNF or Mediator complexes. Combining PRISMA and BioID acts as a powerful strategy to systematically explore the interactomes of intrinsically disordered proteins and their PTM-regulated, multimodal capacity.

**Key points:**

- Integration of proximity labeling and arrayed peptide screen proteomics refines the interactome of C/EBPα isoforms
- Hotspots of protein interactions in C/EBPα mostly occur in conserved short linear motifs
- Interactions of the BAF/SWI-SNF complex with C/EBPα are modulated by arginine methylation and isoform status
- The integrated experimental strategy suits systematic interactome studies of intrinsically disordered proteins

## Introduction

CCAAT enhancer binding protein alpha (C/EBPα) is a lineage instructing pioneering transcription factor involved in cell fate decisions in various cell types, including myeloid cells, adipocytes, hepatocytes, and skin cells. In myelopoiesis, C/EBPα cooperates with other transcription factors and chromatin modifying enzymes to regulate hematopoietic stem cell biology, lineage choice and expression of myeloid genes (Avellino and Delwel, 2017; Zaret and Carroll, 2011). Knockout experiments in mice show that C/EBPα regulates hematopoietic stem cell functions and that its removal blocks progenitor differentiation at the myeloid commitment stage (Bereshchenko et al., 2009; Kirstetter et al., 2008a; Zhang et al., 2004). The intronless *CEBPA* gene is mutated in approximately 10-15% of human acute myeloid leukemia (AML) cases. Mutations located within the N-terminal part of the gene obstruct expression of the full-length isoform p42 C/EBPα and favor expression of the N-terminally truncated isoform p30 C/EBPα (Fasan et al., 2014; Lin et al., 2005; Pabst et al., 2001). Experimental hematology and targeted mouse genetics have demonstrated that p30 *C/EBPα* represents a highly penetrant AML gene (Bereshchenko et al., 2009; Kirstetter et al., 2008b).

Unraveling the protein interaction network of C/EBPα in hematopoietic cells is a prerequisite for the understanding of diverse gene regulatory and epigenetic functions of C/EBPα in normal hematopoiesis and AML. Previous research has shown that the N-terminus of vertebrate C/EBPα harbors short, conserved regions (CRs) that function in a modular and combinatorial fashion to regulate gene expression (Leutz et al., 2011; Tsukada et al., 2011). The N-terminally truncated, leukemogenic p30 C/EBPα isoform contains only one of the transactivation modules (CR1L, TEIII). P30 C/EBPα retains residual gene regulatory and epigenetic capacity to direct myeloid lineage commitment but lacks the cell maturation inducing and proliferation restricting functions of the long C/EBPα isoform p42 (Bereshchenko et al., 2009; Pedersen et al., 2001). Only the C-terminal part of C/EBPα, comprising approximately one quarter of the protein, may adopt a structured basic leucine zipper (bZIP) domain upon binding to DNA target sites (Seldeen et al., 2008). Otherwise, C/EBPα and other family members represent mosaics of intrinsically disordered regions (IDRs) with alternating small linear motifs (SLiM) and molecular recognition features (MoRF), typical hallmarks of gene regulatory proteins that integrate signal transmission, gene regulation, and epigenetic regulation through promiscuous interactomes (Dunker et al., 2002; Dyson and Wright, 2005; Tompa, 2012; Uversky et al., 2005; van der Lee et al., 2014; Ward et al., 2004; Wright and Dyson, 1999). PTMs, downstream of signaling cascades, frequently alter dynamic protein interaction interfaces of IDR/SLiM/MoRF modules and are of high biological relevance but remain difficult to capture with traditional immune affinity-based methods as a result of their transient nature, weak affinity, and/or sub-stoichiometric participation in biomolecular condensates (Perkins et al., 2010; Sabari et al., 2020). Within the C/EBP family, mutation of PTM sites, including phosphorylation (S, T, Y residues), methylation (K, R residues), and acetylation sites (K residues) or deletion of CRs in their N-termini may radically alter their biological functions (Leutz et al., 2011).

Assessment of the known C/EBPα interactome data showed a surprisingly low overlap (less than 5%) between various published immuno-affinity-mass spectrometry (MS) datasets, hampering the deduction of C/EBPα functions and adjunct molecular mechanisms (Cirilli et al., 2017; Giambruno et al., 2013; Grebien et al., 2016). Here, we combined the protein interaction screen using a peptide matrix (PRISMA) method (Dittmar et al., 2019) with biotin proximity labeling identification (BioID) (Roux et al., 2012) to match high confidence interactomes across the primary sequence and PTM sites with the p42/p30 C/EBPα isoforms in living myeloblastic cells. This experimental strategy may be adapted to explore other multi-valent intrinsically disordered transcriptional regulators, many of which are involved in development and disease.

## Results

### Mapping the C/EBPα interactome with PRISMA

We have previously established the PRISMA method to explore the linear interactomes of intrinsically disordered proteins and applied it to C/EBPβ and its PTMs (Dittmar et al., 2019). Here, we slightly modified the PRISMA protocol to explore the linear and PTM-dependent C/EBPα interactome. Briefly, membrane spot-synthesized, tiling peptides that covered the primary sequence of C/EBPα were incubated with nuclear extracts derived from the human promyelocytic leukemia cell line, NB4. Bound proteins of each spot were subsequently analyzed using quantitative mass spectrometry **(Figure 1A)**. The C/EBPα peptide array included 114 peptides with and without PTMs that were designed with a sequence overlap of seven amino acids (**Supplemental Table 1**). PTMs within the bZIP domain were omitted from the PRISMA screen since they may modulate DNA binding in addition to protein interactions. The overall technical reproducibility of the PRISMA screen replicates showed a median Pearson correlation coefficient of 0.85 and discrete patterns of correlation between different C/EBPα regions **(Supplemental Figure 1)**. PRISMA detected a total of 2,274 proteins, of which 785 proteins passed the significance threshold (FDR < 0.1) in at least one peptide spot **(Supplemental Table 2)**. The majority of significant protein interactions were observed in the regions CR2, CR3, CR4 and CR1L, which together correspond to the trans-regulatory regions of C/EBPα (Leutz et al., 2011). Extracted binding profiles, as shown in **Figure 2C**, highlight distinct interaction profiles of selected protein complexes such as Mediator, or BAF/SWI-SNF components, previously shown to interact with C/EBPα.

**Figure 1:**
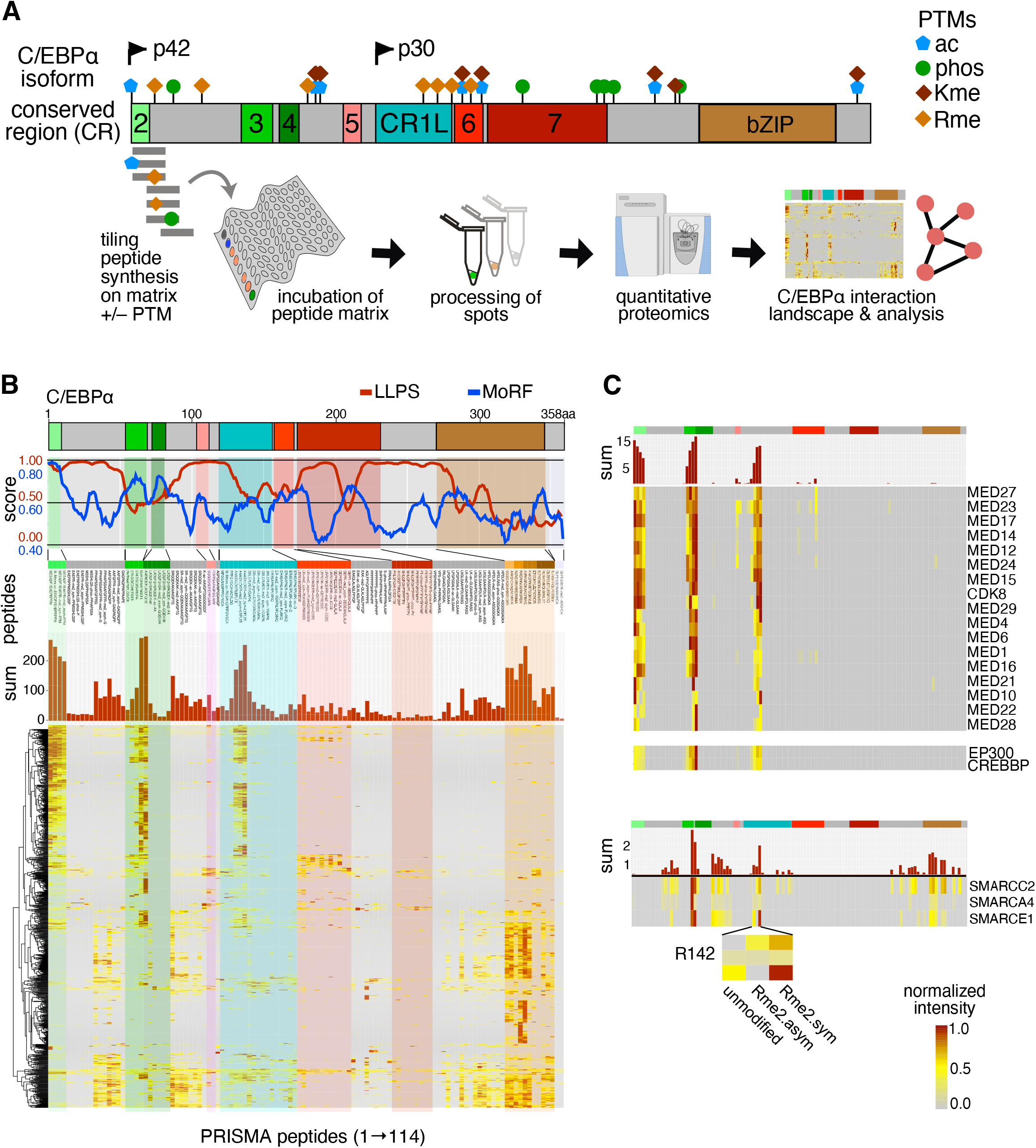
PRISMA delineates the linear interactome of CEBPA across conserved regions and PTM sites. A: Schematic scaled depiction of CEBPA conserved regions (CRs) and post-translational modification (PTM) sites. Spot synthesized, immobilized peptides with and without PTMs covering the complete amino acid sequence of CEBPA were screened for protein interactions with PRISMA. B: Line plots show liquid-liquid phase separation (LLPS; http://protdyn-fuzpred.org) and molecular recognition feature prediction (MoRF; https://morf.msl.ubc.ca/index.xhtml) across the CEBPA sequence. Heatmap shows the binding profile of all significant proteins (y-axis) across CEBPA PRISMA peptides ordered from N-to C-terminus (x-axis). Bar plot on top corresponds to the summed normalized LFQ intensities of all significant proteins in each peptide spot. C: Extracted CEBPA PRISMA profiles of Mediator and P300 complex (top) and BAF/SWI-SNF complex (bottom) subunits.

**Figure 2:**
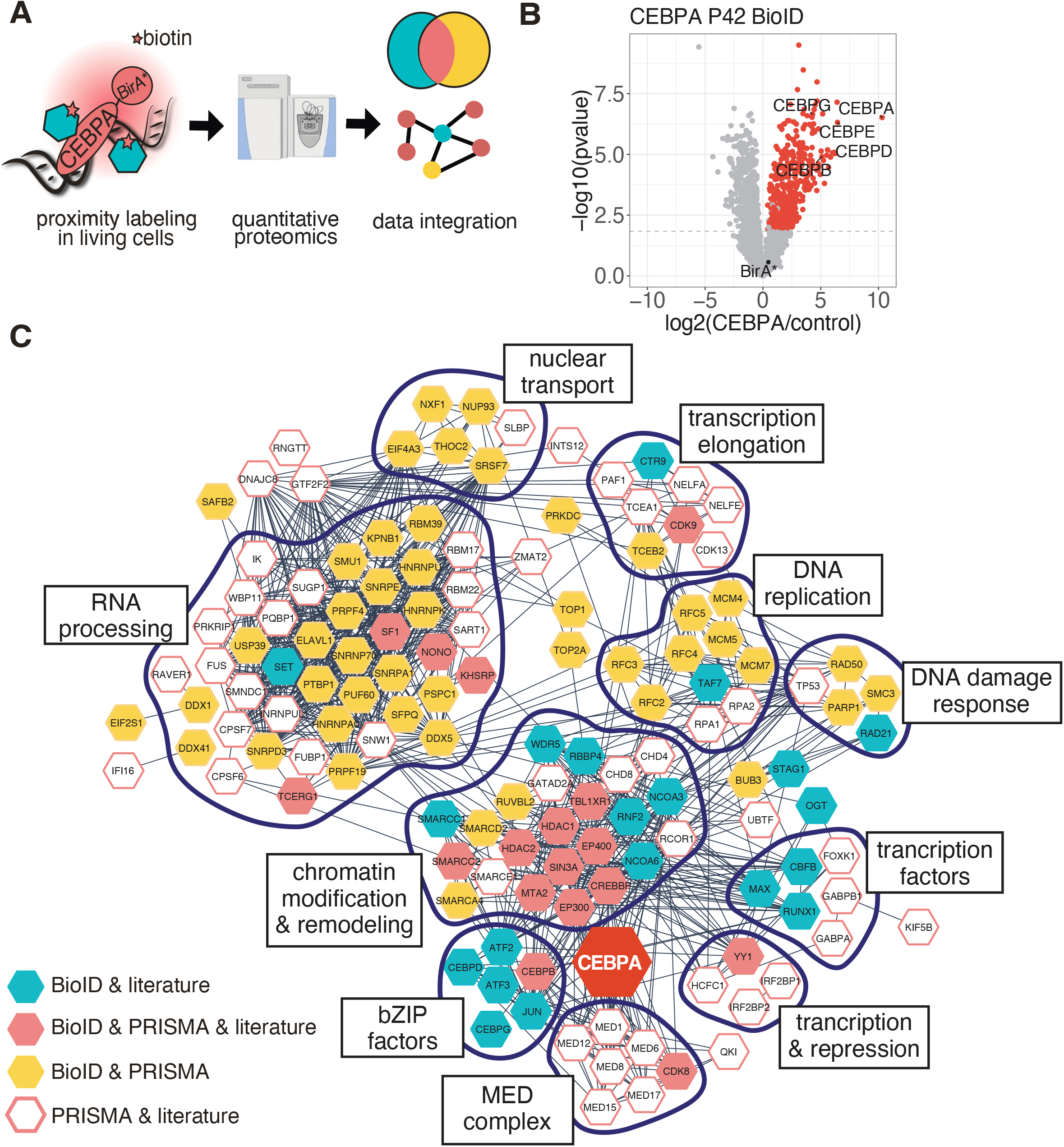
Integration of BioID and PRISMA data revealed a spatial CEBPA interactome with high interconnectivity. A: BioID proximity labelling data obtained in living cells was integrated with biochemical data obtained with PRISMA. B: The CEBPA BioID interactome. Log fold changes (CEBPA/control) of proteins are plotted against their p-values. C: Integrated protein interaction network of CEBPA. Color of nodes corresponds to detection in datasets as depicted in the legend. Edges represent validated protein interactions retrieved from the STRING database. Interactors not connected by any edges were removed from the plot (12 proteins).

The mediator of transcription complex (MED) is an essential transcriptional coactivator in eukaryotes that interacts with RNA Pol II and many transcription factors and co-factors. PRISMA revealed 17 MED proteins with similar binding patterns with dominant peaks in C/EBPα peptides corresponding to CR2,3,4 and CR1L. This finding suggests that the MED-complex interacts with multiple CRs in the N-terminus of p42 C/EBPα and with CR1L/TEIII, which corresponds to the N-terminus of p30 C/EBPα. The acetyltransferases, CBP/p300 (KAT3A/KAT3B), most strongly bound to peptides spanning regions CR3,4 with residual binding in CR2 and CR1L. Interaction with the chromatin remodeling BAF/SWI-SNF complex is essential for the anti-proliferative and differentiation functions of C/EBPα and has been previously found to require CR1L/TEIII in addition to the N-terminus of p42 C/EBPα (Muller et al., 2004; Pedersen et al., 2001). In PRISMA, BAF/SWI-SNF subunits bound to peptides corresponding to CR1L and CR3,4.

C/EBP proteins are extensively decorated with PTMs that may alter their interactome to direct their function (Kowenz-Leutz et al., 2010; Leutz et al., 2011). Of particular interest are a number of C/EBP-family member specific arginine residues, several of which have been shown in C/EBPβ to be targets of mono- and di-methylation, with their mutation profoundly changing C/EBPβ biology (Dittmar et al., 2019; Leutz et al., 2011). We examined C/EBPα for PTMs by mass spectrometry and discovered several known and novel methylation sites, including methylation of R12, R35 and R156 and further identified R142 (CR1L) by a targeted MS-parallel reaction monitoring approach as a novel methylation site **(Supplemental Table 3** and **Supplemental Figure 2)**. The PRISMA data suggested increased binding of the BAF/SWI-SNF subunit SMARCE1 to R142 methylated peptides, as compared to the unmodified counterpart. Other subunits of BAF/SWI-SNF (SMARCA4, SMARCC2) followed a similar trend but their differential binding to the methylated peptide spanning R142 scored somewhat below the threshold set of statistical significance.

Given the sequence conservation between CEBP proteins, we compared the PRISMA derived interactome of C/EBPα with the previously published C/EBPβ PRISMA interactome (Dittmar et al., 2019). Despite some sequence differences in homologous regions and peptides, several shared interactors were identified. For example, the mediator complex was found to bind to the same homologue CRs in both proteins **(Supplemental Figure 3)**, confirming the functional similarity of these regions within the CEBP family (Jones et al., 2002).

**Figure 3:**
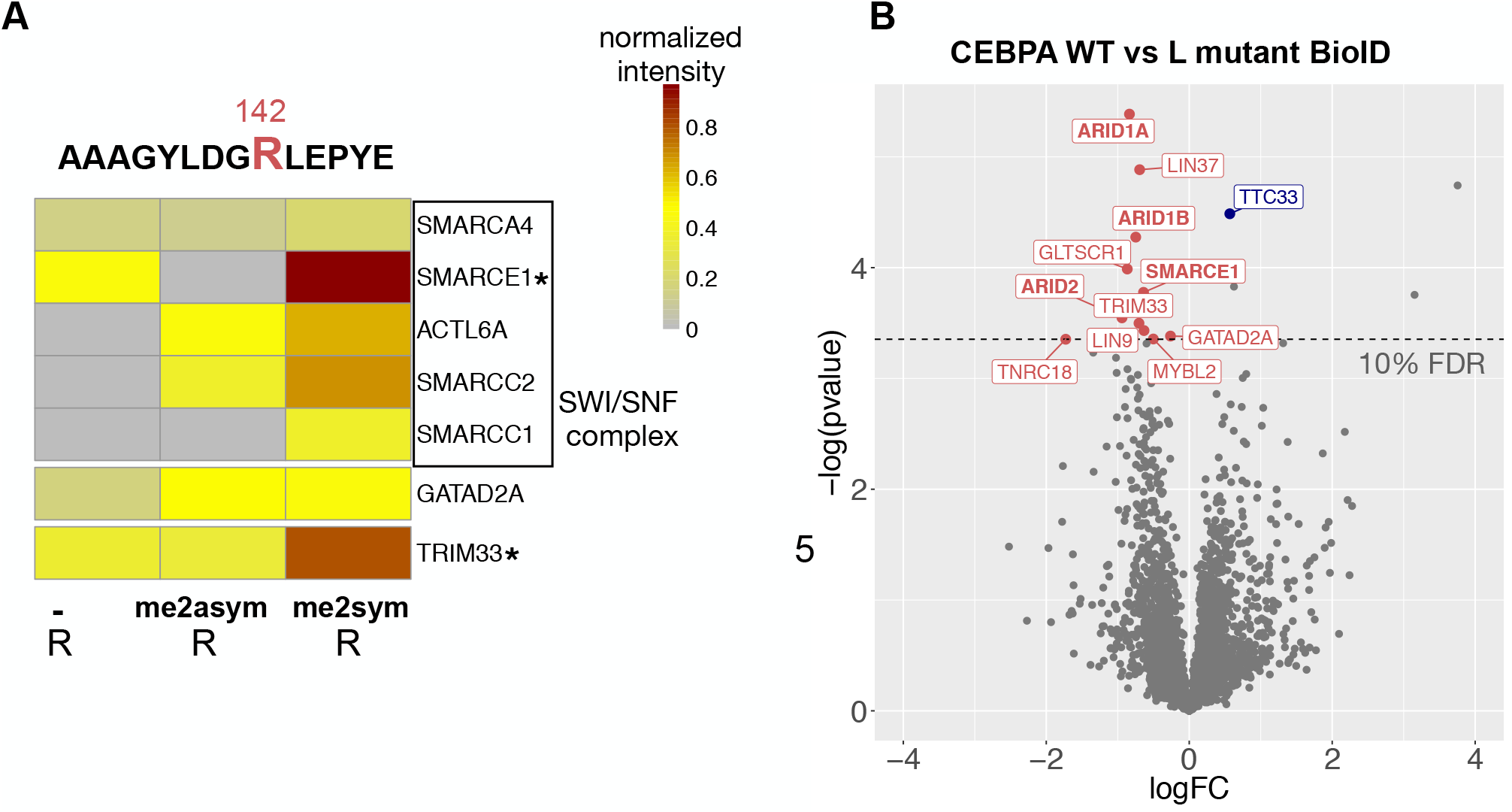
BAF/SWI-SNF complex subunit SMARCE1 preferentially interacts with R142 methylated CEBPA. A: PRISMA interaction profile of proteins differentially binding to CEBPA peptides centered at R142. Peptide sequence is shown on top and methylation status of R142 is shown at the bottom. Stars denote significance (< 0.1 FDR) in R142 dimethylated peptide B: BioID experiments with wildtype CEBPA and trR>L CEBPA. Proteins passing the significance threshold in wildtype to mutant comparison and significant compared to controls are marked in color. Significant BAF/SWI-SNF subunits are indicated by bold letters.

### Cross-validation of PRISMA and BioID-C/EBPα interactomes

Next, we compared the interactomes derived from PRISMA and BioID proximity labeling data to generate a myeloid live cell validated linear C/EBPα interactome (**Figure 2 A**). Briefly, NB4 cells were transduced with a Tet-On inducible lentiviral construct encoding a promiscuous biotin ligase (BirA*) fused to the C-terminus of C/EBPα. As controls, we used non-induced NB4 C/EBPα-BirA* cells or cells expressing only the ligase moiety (NB4 BirA*). Proximity labeling identified 397 C/EBPα proximal interactors (FDR < 0.05). Among the most enriched proteins were several transcription factors of the C/EBP and ATF families, representing known heterodimerization partners of C/EBPα (Tsukada et al., 2011) and thus confirming successful BioID labeling (**Figure 2B, Supplemental Table 4**).

In total, 88 proteins overlapped between the PRISMA and BioID derived C/EBPα interactomes of which 21 were previously identified interactors, including members of chromatin remodeling and histone acetylation / deacetylation complexes. In addition, 49 proteins significant in PRISMA and 26 proteins significant in BioID have been previously described to interact with C/EBPα (Chatr-Aryamontri et al., 2017; Grebien et al., 2016; Szklarczyk et al., 2015). Taken together, 137 proteins represent the subset of myeloid C/EBPα interactors with the highest confidence that can be depicted across the linear C/EBPα sequence +/- PTM sites. These interactors show high connectivity according to experimentally validated interactions listed in the STRING database **(Figure 2C)**. Using the spatial information provided by PRISMA, we could also reveal distinct functional roles of individual CRs including chromatin remodeling (CR3,4; CR1L; bZIP), transcriptional regulation (CR2; CR3,4; CR1L; CR6; bZIP) and hematopoietic progenitor cell differentiation (CR3,4) (**Supplemental Figure 4)**. CR3,4 is of particular interest since it contains most of the significantly enriched GO terms, confirming the importance of the core transactivating region of p42 C/EBPα. Although several proteins interacted with CR7, no GO terms were enriched with this region, pointing towards functional heterogeneity.

**Figure 4:**
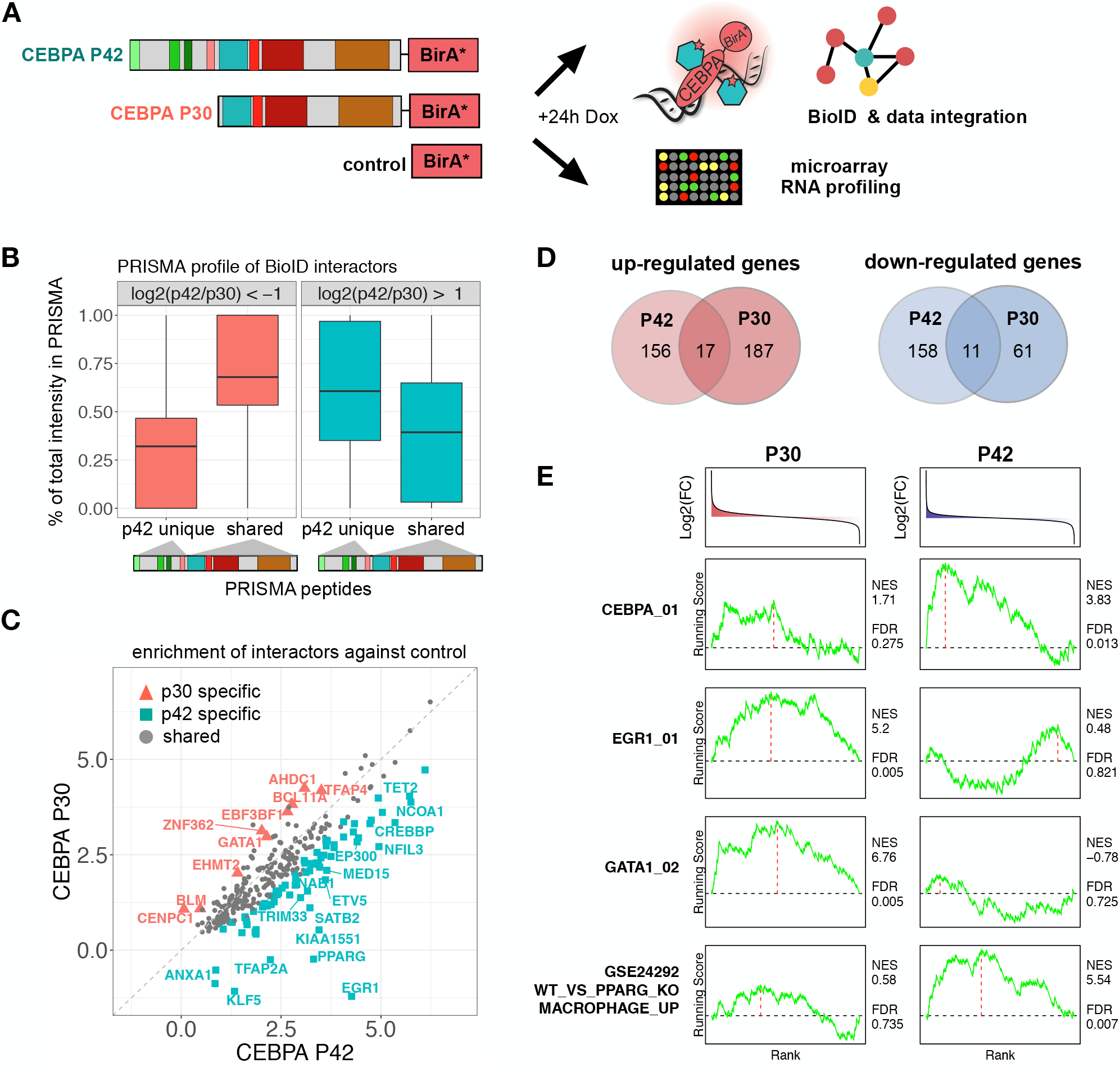
BioID with CEBPA p42/p30 isoforms disclose isoform-specific interactors in conjunction with PRISMA profiles. A: BioID interaction proteomics and microarray RNA expression profiling were performed with CEBPA isoform (p42 or p30) expressing cells. B: PRISMA profiles of interactors showing preference for p42 (log2(p42/p30) > 1) or p30 (log2(p42/p30) < − 1) in BioID. LFQ intensities of proteins were summed across PRISMA peptides corresponding to exclusively p42 or both isoforms. The resulting numbers were divided by the total LFQ intensity of each protein across all PRISMA peptides. PTM-modified peptides were omitted from the calculations. C: Relative enrichment of CEBPA interactors against the control, p42 is plotted against p30. Proteins marked in color passed 0.1 FDR threshold in a direct comparison. D: Overlap of up- and downregulated genes as detected by microarray gene expression profiling (comparison to BirA* cells, FDR < 0.05, absolute fold change(FC) > 1). E: Induced gene expression changes were subjected to ssGSEA analysis. Normalized enrichment score (NES) and FDR of informative gene sets are displayed next to running score line plots. The running score was calculated with Kolmogorov-Smirnov (K-S) running sum statistics.

### Arginine methylation-dependent interaction of BAF/SWI-SNF complex subunits

Differential interactions with post-translationally modified peptides in the primary C/EBPα sequence were also detected by PRISMA. Among them was the BAF/SWI-SNF complex that has been previously described to interact with C/EBPα (Pedersen et al., 2001). PRISMA revealed that the interaction with the SMARCE1 component is modulated by the newly discovered arginine methylation site R142 within CR1L/TEIII **(Figure 3A)**. Accordingly, we performed BioID with a R142/149/156 to L142/149/156 mutant (triple R to L mutation; tR>L) of C/EBPα to examine the PTM-dependent BAF/SWI-SNF interaction in more detail. In accordance with the PRISMA results, SMARCE1 and three additional BAF/SWI-SNF subunits (ARID1A, ARID1B, ARID2) were significantly enriched in the p30 C/EBPα-tR>L mutant, as compared to the WT p30 C/EBPα-BioID **(Figure 3B)**. BioID with p30 C/EBPα-tR>L also verified the methylation-dependent interaction with the E3 ubiquitin ligase TRIM33 and the NuRD complex component GATAD2A, as also detected by PRISMA. In addition, several Myb-Muvb/DREAM complex members (LIN9, LIN37, MYBL2) were identified as p30 C/EBPα-tR>L-specific interactors but were not detected in PRISMA.

### C/EBPα isoform specific interactions detected with BioID

The PRISMA data predicted that p30 C/EBPα isoform can still function to recruit major components of the transcriptional and epigenetic machinery, albeit with lower efficiency compared to p42 C/EBPα. To further examine the differences between the two C/EBPα isoforms, we expressed p42 and p30 C/EBPα as BioID fusions in NB4 cells to compare their BioID interactomes. The isoform-specific interactomes **(Figure 4A, Supplemental Table 4)** confirmed the binding profiles observed in PRISMA: p42 specific interactors bound more strongly to PRISMA peptides corresponding to the N-terminal, p42 unique part of C/EBPα, while p30 specific interactors predominantly bound to PRISMA peptides corresponding to C-terminal C/EBPα regions **(Figure 4B)**. In accordance with the PRISMA interactions, most interactors identified by in vivo proximity labelling were found to interact with both C/EBPα isoforms (**Figure 4C**). These data suggest that multvalent interactions with different affinities may occur with distinct C/EBPα CRs, including CR1L as part of p30 C/EBPα. Overall, p42 C/EBPα-BioID and p30 C/EBPα-BioID pulldowns revealed 71 and 9 interactors that preferentially interacted with p42 and p30 C/EBPα, respectively.

Proteins differentially interacting with the p30 C/EBPα isoform and its PTMs may pose a selective vulnerability for AML cells expressing p30 C/EBPα. We therefore extracted CRISPR Cas9 knockout study derived dependency scores of the nine p30 C/EBPα specific interactors in 18 different AML cell lines using the DepMap Portal **(Supplemental Figure 5)** (Tsherniak et al., 2017). Among the AML cell lines inspected, 9 out of 18 scored as sensitive to TFAP4 knockout (threshold < – 0.5). Two and one cell line tested were sensitive to GATA1 and BCL11A or BLM knockout, respectively. The p30 C/EBPα specific binding of the BAF/SWI-SNF complex member BCL11A further highlights context-specific regulation of BAF/SWI-SNF complex interaction to C/EBPα isoforms.

To analyze connections between the C/EBPα isoform specific interactome with the transcriptome we also performed RNA expression analysis of NB4 cell lines ectopically expressing p42 or p30 C/EBPα. We found that despite the largely overlapping interactome, the two isoforms induced differential gene expression changes (**Figure 4D**) that could be attributed to differential interactions with other transcription factors. Gene expression profiles were analyzed using gene set enrichment analysis (GSEA) employing immunologic and transcription factor target databases from the molecular signature database (Subramanian et al., 2005) (**Figure 4E**). The erythroid transcription factor GATA1 was found to interact differentially with p30 C/EBPα; GSEA analysis detected enrichment of a GATA1 signature specifically in p30 C/EBPα expressing cells. Likewise, PPARG specifically interacts with p42 C/EBPα and p42 expressing cells also displayed significant enrichment of a previously published macrophage PPARG gene signature (Roszer et al., 2011). Data from BioID experiments indicated that EGR1 specifically interacted with p42 C/EBPα. Interestingly, we found that a gene signature based on the presence of EGR1 motifs was enriched in p30 but not p42 C/EBPα expressing cells. RNA expression levels of EGR1 were upregulated by both p42 and p30 expression. Indirect interaction of EGR1 with DNA not at EGR1 motifs, but through other CEBPs, has been described previously (Jakobsen et al., 2013) and may contribute to the differential EGR1 signature observed. Taken together, our data suggest that the specific interactions of C/EBPα isoforms with lineage defining transcription factors are implicated in co-regulation of target genes in the hematopoietic system.

## Discussion

In the present study, we combined peptide array screening and BioID to fine map the interactome of the intrinsically disordered transcription factor C/EBPα. The integration of both methods overcame limitations of classical immuno-affinity based interactomes of intrinsically disordered proteins. Contribution of individual PTMs and protein regions to the interactome remain difficult to deconvolute in classical pull-down MS approaches, while PRISMA reveals both SLiM- and PTM-dependent interactions (Dittmar et al., 2019; Meyer and Selbach, 2020). Moreover, transient interactions mediated by IDRs/SLiMs are easily lost during affinity-purification. In contrast, BioID covalently biotinylates proteins that engage with C/EBPα in the living cell and thus enables the detection of transient and dynamic interactions (Roux et al., 2012). Altogether, PRISMA and BioID revealed 785 and 397 interacting proteins, respectively, and the overlay of C/EBPα BioID and PRISMA data sets with public resources disclosed a linear, high confidence C/EBPα core-interactome of 137 proteins. The C/EBPα core-interactome can be depicted as an interaction landscape across the C/EBPα sequence and PTM sites and its components were found to be highly interconnected. In addition to known C/EBPα interactors that operationally represent positive controls, we also discovered novel associations of C/EBPα with proteins involved in gene expression and epigenetic regulation, chromosome organization, RNA processing, and DNA replication, extending the repertoire of myeloid C/EBPα targets and associated functions. The majority of C/EBPα interactions were nevertheless detected in only one of the two datasets. Some of the unique C/EBPα interactors detected by BioID may require more complex, simultaneously occurring, multi-site interactions or multiple induced fit processes and were missed in PRISMA. Discrepancies may also relate to restriction of BioID to proximal interactions, the absence of suitable biotinylation sites, or stability during the labelling period. PRISMA, on the other hand, may readily detect distal, secondary interactors, such as in large protein complexes.

PRISMA revealed major hotspots of SLiM-based protein interactions in C/EBPα that strongly correlated with data derived from the related transcription factor C/EBPβ (Dittmar et al., 2019; Leutz et al., 2011). The interactors shared between C/EBPα and C/EBPβ may help to explain the partially redundant functions of both proteins (Chen et al., 2000; Hirai et al., 2006; Jones et al., 2002). Many of the co-regulators identified also displayed multiple interactions with several SLiMs/MoRFs/CRs, predominantly in CR2,3,4 and CR1L. These regions correspond to previously defined trans-regulatory regions found within the C/EBP family (Leutz et al., 2011). The same conserved regions scored highly as molecular recognition features (MoRF; **Figure 1B**) that may undergo induced folding and transient disorder-to-order transition during contact with partner proteins (Oldfield et al., 2005). For example, the region CR3,4 of C/EBPε was previously shown to fold into two short orthogonal amphipathic helical regions on interaction with the TAZ2 domain of CBP (Bhaumik et al., 2014). Likewise, CBP/p300 was found to interact with PRISMA peptides covering CR3,4 of C/EBPα, in concordance with homologous regions in C/EBPβ (Dittmar et al., 2019). Interestingly, CR3,4 in C/EBPα and C/EBPε are separated by only few amino acids, whereas species-specific IDRs separate both CRs in C/EBPβ, suggesting structural spatial flexibility and independent, yet combinatorial functions of CR3,4.

Many regulatory proteins involved in signaling, gene expression, and epigenetics harbor IDRs and PTMs that determine their function and connectivity (Dyson and Wright, 2005, 2016). Although protein interactions directed by SLiMs are in general of low affinity, they may nevertheless exhibit high specificity. This raises the question of how specificity is achieved. Specificity may be forged by the combined action of several SLiMs and PTMs to regulate selective interactions of a subset of SLiMs at a time. Multivalent, promiscuous interactions of transcription factors with co-regulatory proteins have been observed before and may relate to context dependent contacts during dynamic gene regulatory processes (Brzovic et al., 2011; Clark et al., 2018; Vojnic et al., 2011). IDRs within and between SLiMs permit structural flexibility and dynamic interactions, which can also drive the formation of specific protein condensates with transiently favorable interactions (Hahn, 2018). This concept is in accordance with the promiscuous nature of SLiMs and may reflect dynamic multi-modal regulation with the rapid exchange of alternative interaction partners (Dyson and Wright, 2016; Ivarsson and Jemth, 2019). Modular transactivation domains in conjunction with multi-valency are thought to participate in or to initiate biomolecular condensates that involve dynamic, “fuzzy” interactions with multiple co-regulators in interaction hubs (Boija et al., 2018; Brzovic et al., 2011; Chong et al., 2018; Hahn, 2018; Martin and Holehouse, 2020; Tuttle et al., 2018). Remarkably, the C/EBPα primary sequence shows a very high degree of “fuzziness” with strong predictions as initiator regions of liquid-liquid phase separation in the N-terminus **(Figure 1B, red line)**, while the PRISMA interaction hotspots largely coincide within regions that show maxima of MoRF predictions **(Figure 1B, blue line)** (Horvath et al., 2020; Malhis et al., 2016).

Our data show that PTMs in C/EBPα may orchestrate multi-modal functions of C/EBPα by modulating interactions with the components of transcriptional and epigenetic machinery. The comparison of C/EBPα WT and the R142L methylation mimicry mutant by proximity labeling confirmed the differential interaction with BAF/SWI-SNF components, as detected by PRISMA. These data are consistent with previous findings showing that BAF/SWI-SNF interacts with CR1L (Muller et al., 2004; Pedersen et al., 2001). The newly discovered R142 methylation-dependent differential interaction of BAF/SWI-SNF will help to conceive hypothesis driven approaches to explore arginine methyltransferase-dependent epigenetic downstream events.

Data presented here further demonstrate that most of the C/EBPα interactors bind to both C/EBPα isoforms, whereas a subset of 71 and 9 proteins preferentially interacted with the p42 and p30 C/EBPα isoform, respectively. Many of the shared C/EBPα interactors showed diminished signal strength in p30 C/EBPα BioID, supporting the concept of p30 as a “weak” C/EBPα variant with relatively few exclusive binding partners. This view is supported by the fact that defective myelopoiesis in the absence of C/EBPα was largely rescued by p30 C/EBPα, although p30, in the absence of p42 C/EBPα, eventually elicits AML (Bereshchenko et al., 2009; Kirstetter et al., 2008). Interestingly, several of the p30 C/EBPα-specific binders (TFAP4, GATA1, BCL11A) affect the survival of AML cell lines. The p30 C/EBPα specific interaction with GATA1 may further hint at a role at the branch point of erythroid / myeloid commitment in replicative multi-potential progenitors (Drissen et al., 2019), whereas p42 C/EBPα specific interactors, such as EGR1 or PPARγ, are important regulators of myeloid differentiation (Chinetti et al., 1998; Krishnaraju et al., 2001; Lefterova et al., 2010; Lefterova et al., 2008; Mildner et al., 2017; Nguyen et al., 1993; Roszer et al., 2011; Tontonoz et al., 1994).

In summary, the cross-validated myeloid C/EBPα interaction map presented in this study may serve as a resource for further exploration of the biological importance of individual and combinatorial SLiM and PTM functions of C/EBPα. Beyond the C/EBPα,β interactomes, the integration of PRISMA and BioID approaches may help to explore the linear and PTM dependent interactomes of many other important intrinsically disordered proteins involved in cell signaling and cell fate determination.

## Materials and Methods

### Cell culture

NB4 cells were acquired from Leibniz Institute DSMZ-German Collection of Microorganisms and Cell Culture, Germany (DSMZ no.: ACC 207). Cells were cultivated in a humidified incubator at 37°C, 5% CO2 in RPMI1640. For metabolic labeling, NB4 cells were grown in SILAC RPMI1640 supplemented with 10% dialyzed FCS, 100 mg/ml penicillin-streptomycin, 25mM HEPES, 28 µg/ml L-arginine and 48.67 µg/ml L-lysine (light) 13C615N2 (heavy lysine) or L-lysine D4 (medium heavy lysine).

### Plasmids and Generation of Cell Lines

The rat C/EBPα p30 3L mutant (R140; 147; 154 mutated to leucine) was generated using the QuickChange Site Directed Mutagenesis Kit according to the manufacturer’s protocol (Agilent #200519). The BirA Ligase containing plasmid was purchased (Addgene #64395). The BirA Ligase was cloned in frame to the C/EBPα C-terminus using BamHI and XhoI restriction sites. PCR primer: 5’-CGCGGATCCAGCGGTGGAAGTGGTGGCCTGAAGGACAACACCGTG and 3’-TGCTCTAGACTCGAGTTATTTATCGTC. The C/EBPα p42, p30 or p30 R3L-BirA Ligase fragments were cloned into the pENTRY2B vector (Clontech #3064) using BamHI and XhoI restriction sites and subsequently introduced into the pInducer21 GFP lentiviral vector (Clontech #3044) using the Gateway LR Clonase TM II cloning kit (ThermoFisher Scientific #11791-020). Viral supernatants were obtained from Lenti-X 293T cells (Clontech #632180) transfected with either pInducer21 GFP lentiviral vector or pInducer21 GFP constructs containing either p42, p30 WT-BirA Ligase, p30 R3L-BirA Ligase, or pInducer21 GFP-BirA ligase only using Lenti-X Packing Single Shots (Clontech #631275) according to the manufacturer’s protocol. NB4 cells were centrifuged (1 h, 900g) with infectious supernatant collected 72 h after transfection and 8 µg/mL hexadimethrine bromide and left for recovery overnight. GFP positive NB4 cells were sorted twice using an Aria II sorter (Becton Dickinson) three days after infection.

### NB4 induction

NB4 cells were briefly treated with differentiation inducing agents prior to harvesting. SILAC labeled NB4 cells were seeded during exponential growth in SILAC media supplemented with 2µM all-trans-retinoic acid (ATRA; heavy), 50nM Tetradecanoylphorbol-acetat (TPA; medium heavy) or solvent only (light). Cells were harvested after 12h (ATRA and solvent only treated cells) or 6h (TPA treated cells) and nuclear extracts were prepared as described.

### Nuclear extract preparation

Nuclear extracts from NB4 cells were prepared as described previously (Dignam et al., 1983) with slight modifications. NB4 cells were harvested by centrifugation at 1000g, 4 min at 4°C and washed twice with ice cold PBS. Packed cell volume (pcv) was estimated and cells were resuspended in 5xpcv of ice-cold hypotonic buffer (10mM HEPES pH 7.5, 10mM NaCl, 3mM MgCl2) supplemented with protease inhibitors. Cells were incubated on ice for 5 min, followed by addition of dodecyl-β-D-maltosid (DDM) to a final concentration of 0.05%. The sample was vortexed briefly and immediately centrifuged for 5 min at 600g, 4°C. The cytosolic fraction was removed and the nuclei were washed with 20xpcv hypotonic buffer (5 min, 600g, 4°C). The supernatant was removed and the nuclei were washed with 20xpcv PBS (5 min, 600g, 4°C). The nuclei were extracted with 2/3xpcv of high salt buffer (20Mm HEPES pH 7.5, 400mM NaCl, 1mM EDTA ph 8, 1mM EGTA pH 8, 20% glycerol, 1mM DTT) supplemented with protease inhibitors while shaking on a tubeshaker at 4°C for 20 min at 750 rpm. Nuclear extracts were cleared by centrifugation at 18000g for 20 min at 4°C and the buffer was exchanged to membrane binding buffer (20mM HEPES pH 7.5, 400mM NaCl, 1mM EDTA ph 8, 1mM EGTA pH 8, 25% glycerol, 1mM DTT) by gel filtration with PD MidiTrap G10 columns (GE healthcare) according to instructions of manufacturers.

### Protein Interaction Screen on a peptide Matrix

Custom PepSpot cellulose membranes were ordered from JPT and PRISMA was performed as described before (Dittmar et al., 2019) with slight adaptations. All washing and incubation steps were performed at 4°C on a rocking platform set to 700 rpm. Membranes were wetted in membrane binding buffer for 15 min, followed by a blocking step with 1 mg/ml yeast tRNA in membrane binding buffer for 10 min. Membranes were washed 5 x for 5 min with membrane binding buffer and incubated with nuclear extracts on ice for 30 min. The protein extract was removed and the membranes were washed 3 x 5 min with membrane binding buffer. The individual peptide spots were punched out with a 2mm biopsy puncher and placed into single wells of a 96 well plate containing 20µl denaturation buffer (6M urea, 2M thiourea, 10mM HEPES pH 8). The samples were digested in solution on a PAL robot system. In brief, proteins were reduced with 1mM TCEP for 30 min followed by alkylation with 5mM CAA for 20 min. To each sample 0.5 µg of sequencing grade lysyl endopeptidase (LysC) was added. Samples were digested for 2h before being diluted with four volumes of 50mM ammonium-bi-carbonate and continuation of the digestion over night at room temperature. Digested samples were acidified with TFA and desalted with C18 stage tips as described before (Rappsilber et al., 2003).

### LC MS/MS

Desalted and dried peptides were resuspended in MS sample buffer (3% ACN/ 0.1% FA) and separated online with a Easy-nLC™ 1200 coupled to a Q-exactive+ or a Q exactive HF-X mass spectrometer equipped with an orbitrap electrospray ion source. Samples were separated on line on a 20 cm reverse-phase column (inner diameter 75µm) packed in house with 3 µm C18-Reprosil beads with a linear gradient ramping from 3% to 76% acetonitrile. PRISMA samples were separated with a 1h gradient and MS data was acquired on a Q-exactive+ in data dependent acquisition mode with a top10 method. Full scan MS spectra were acquired at a resolution of 70 000 in the scan range from 300 to 1700 m/z, automated gain control (AGC) target value of 1e^6^ and maximum injection time of 120ms. MS/MS spectra were acquired at a resolution of 17500, AGC target of 1e^5^ and maximum IT of 60 ms. Ions were isolated with a 2 m/z isolation window and normalized collision energy was set to 26. Unassigned charge states and single charged precursors were excluded from fragmentation and dynamic exclusion was set to 20s. BioID pulldowns were separated on with a 2h gradient and MS data was acquired on a Q-exactive HF-X in data dependent acquisition mode with a top20 method. Full scan MS spectra were acquired at a resolution of 60000 in the scan range from 350 to 1700 m/z, automated gain control (AGC) target value was set to 3e^6^ and maximum injection time to 10ms. MS/MS spectra were acquired at a resolution of 30000, AGC target of 1e^5^ and maximum IT of 86 ms. Ions were isolated with a 1.6 m/z isolation window and normalized collision energy was set to 26. Unassigned charge states and ions with a chare state of one, seven or higher were excluded from fragmentation and dynamic exclusion was set to 30s. The mass spectrometry proteomics data and search results have been deposited to the ProteomeXchange Consortium via the PRIDE partner repository (Perez-Riverol et al., 2019) with the dataset identifier PXD022903.

### BioID experiments

NB4 stable cell lines were grown in RPMI supplemented with tetracycline free FCS. Cells were seeded in exponential growth phase in media supplemented with 1mM biotin and 1µg/ml doxycycline. Cells were harvested after 24h by centrifugation and washed twice with ice cold PBS. Cell pellets were resuspended in modified RIPA buffer (lysis buffer: 50 mM tris-HCl (pH 7.2), 150 mM, NaCl, 1% NP-40, 1 mM EDTA, 1 mM EGTA, 0.1% SDS, 1% sodium deoxycholate, freshly added protease inhibitors) and incubated on ice for 20 min. Samples were sonicated with a probe sonicator for 5 pulses and centrifuged for 20 min at 4°C, 20 000g. For each pulldown (1x 15 cm dish), 80µl neutravidin-agarose bead slurry (Thermo Fisher Scientific) was used. Beads were washed twice in lysis buffer and added to the protein extracts. The samples incubated rotating at 4°C for 2.5 h. Beads were washed 3x with lysis buffer, 1x with 1M KCl, 1x 2M Urea in 50mM Tris pH8 and 3x with 50mM Tris pH8. Washing buffers were kept on ice and each washing step was performed with 1ml, inverting the tube 5 times and then centrifuging for 1 min at 2000g to pellet the beads. Washed beads were resuspended in 80µl urea/trypsin buffer (2M urea, 50mM Tris pH7.5, 1mM DTT, 5µg/ml trypsin) and incubated 1h at RT on a tubeshaker at 1000rpm. The supernatant was transferred into a fresh tube, the beads washed twice with 60µl 2M urea/50mM Tris pH7.5 and the supernatant combined with the supernatant from the previous step. Residual beads were removed by an additional centrifugation (1 min, 5000g) and sample transfer step. Eluted proteins were reduced with 4mM DTT for 30 min on a tubeshaker (RT, 1000 rpm). Proteins were alkylated with 10mM IAA in the dark for 45 min (RT, 1000 rpm) and digested over night with 0.5µg trypsin at RT on a thermoshaker at 700 rpm. For an AspN digest, 0.5 µg of trypsin and 0.5 µg AspN were added to the sample. Following overnight digestion, samples were acidified by adding TFA and desalted with stage tips.

### Targeted MS analysis of R142 methylation

Synthetic heavy peptide standards with the sequence DGRLEPLEYER and DGR[me]LEPLEYER were custom synthesized by JPT (spiketides L, labeled at the C-terminus with heavy arginine (Arg10)). CEBPA BioID pulldowns were digested with AspN and trypsin and desalted as described. Samples were resuspended in MS sample buffer containing 100 fmol/µl of synthetic heavy peptides. Parallel reaction monitoring (PRM) measurements were acquired on a Q exactive HF-X mass spectrometer coupled to an Easy-nLC™ 1200 HPLC system. Peptides were separated on a 60 min gradient ramping from 2% to 76% acetonitrile. Data acquisition mode cycled between a Top1 MS/data dependent MS2 and data independent measurement of an inclusion list that included the m/z of the synthetic heavy peptides as well as their light counterparts coming from the sample. Resolution of MS2 for data independent acquisition was 60 000 with an AGC target of 1e6, 200 ms maximum IT, 0.7 m/z isolation window and NCE of 27. PRM data was analyzed with Skyline (Pino et al., 2020). Identity of peptides was verified by comparison with the elution profile and fragmentation spectrum of heavy peptide standards.

### Mass spec raw data processing with MaxQuant

Mass spectrometry raw files were processed with MaxQuant (version 1.5.2.8) (Cox and Mann, 2008) searching against a human protein database containing isoforms downloaded from uniprot (June 2017) and a database including common contaminants. Fixed modifications were set to carbamidomethylation of cysteines and variable modifications were set to methionine oxidation and N-terminal acetylation. For BioID experiments, lysine biotinylation was added as an additional variable modification. Depending on digestion mode (trypsin or LysC only), enzyme specificity was selected with a maximum of 2 missed cleavages per peptide. The initial allowed mass deviation of the precursor ion was up to 6 ppm and 20 ppm for fragments. False-discovery rate was set to 1% on protein and peptide level. For SILAC measurements, the requantify option was enabled and minimum ratio count was set to 2. For LFQ analysis, the match between run and fast LFQ option was used with default settings.

### Data analysis of mass spec data

Statistical analysis of the dataset was performed using R-statistical software package (version 3.4.1). The protein groups output file from MaxQuant was filtered for contaminants, reverse hits and proteins only identified by site. Minimum peptide count for SILAC and LFQ data was at least 2 peptides per protein group. For LFQ datasets, proteins were filtered by detection in at least 3 replicates of the same sample and missing values were imputed from a distribution at the detection limit of the mass spectrometer. For this purpose a normal distribution was created for each run, with a mean 1.8 standard deviations away from the observed mean and a standard deviation of 0.3 x the observed standard deviation. LFQ data was analysed with a two sample moderated t-test (Limma package) and p-values were corrected for multiple testing with Benjamini-Hochberg procedure. The significance cutoff of CEBPA interactors was enrichment against both controls (BirA* only and cells not treated with doxycycline) with an FDR < 0.05. For isoform and tR>L mutant specific interactors, threshold was enrichment against both controls with an FDR < 0.05 and FDR < 0.1 in the pairwise comparison of isoforms and mutants. Interactors overlapping between PRISMA and BioID or BioID and literature (STRING and BioGrid database, Grebien et al., Giambruno et al.) were visualized with the STRING app built into cytoscape (v3.71) (Shannon et al., 2003). Known interaction (experimentally validated or deposited in databases with a score > 0.5) were visualized as edges. Interactors not connected by any edges were removed from figure 2C (12 proteins). GO term analysis of interactors was performed with DAVID functional annotation tool (version 6.8) (Huang et al., 2009). PRISMA data was analysed with a two sample moderated t-test (Limma package), creating a specific control group for each peptide that contained all other peptides excluding peptides with a sequence overlap ≥ 50%. P-values were corrected for multiple testing with Benjamini-Hochberg procedure, significance cut off was < 0.1 FDR. To obtain binding profiles of interactors, LFQ intensities of significant proteins were normalized between 0 and 1 across all PRISMA peptides.

### Microarray

Cells were harvested at exponential growth phase and seeded at 0.5 × 10^6^ cells/ml in RPMI supplemented with or without 1µg/µl doxycycline as independent biological triplicates. After 24h cells were harvested via centrifugation (1200 g, 5 min, 4°C) and washed once with 1x ice cold PBS. Total RNA was extracted with the RNeasy Mini Kit (Qiagen) following manufacturer’s instructions. DNA was removed with the DNase Max Kit (Qiagen) following manufacturer’s instructions. RNA integrity was assessed using a Fragment Analyzer system and a standard sensitivity RNA kit. All RQN scores were above 9.1. RNA expression analysis was performed with Affymetrix Human ClariomS® microarray using the WT Plus Reagent kit (ThermoFisher Scientific). As starting material, 100ng of total RNA was used and RNA was prepared for hybridization with the GeneChip® Whole Transcript (WT) PLUS Reagent Kit (ThermoFisher Scientific) following manufactures instructions. CEL files were processed using the standard Transcriptome Analysis Console (TAC 4.0) software. Expression values were automatically normalized and summarized using SST-RMA method. Only mRNAs with log2 expression above 6 in at least one sample were considered for further analysis. Statistical analysis was performed using LIMMA package of R/Bioconductor and p-values were adjusted using Benjamini-Hochberg’s FDR. Microarray data have been deposited in the ArrayExpress database at EMBL-EBI (www.ebi.ac.uk/arrayexpress) under accession number E-MTAB-9947.

### GSEA of microarray data

Gene expression changes induced by CEBPA isoforms were calculated by comparing microarray data from NB4 cells expressing CEBPA (P42 or P30) against BirA* expressing cells. Averaged log2 fold changes were used as input for the single sample gene set enrichment analysis (ssGSEA) analysis tool (Barbie et al., 2009) implemented in R (https://github.com/broadinstitute/ssGSEA2.0) using standard parameters. Data was analyzed employing immunologic and transcription factor target gene sets from the molecular signature database (Subramanian et al., 2005). The running score displayed in Fig.4 was calculated with Kolmogorov-Smirnov running sum statistics.

## Author contributions

Conceptualization, A.L., G.D., E.R., D.P.; Methodology, G.D., A.L., E.R., E.K.L., V.S.; Investigation and Validation, E.R., V.S., E.K.L.; Resources, G.D., A.L., P.M., U.R., N.N., P.N.; Data curation, E.R., K.Z., A.L.; Writing original draft, E.R.; Writing, review and editing, E.R., A.L., E.K.L., P.M. G.D.; Visualization, E.R., A.L.; Supervision, project administration, and funding acquisition, A.L. (ORCID: 0000-0001-8259-927X), G.D. (ORCID: 0000-0003-3647-8623), P.M. (ORCID: 0000-0002-2245-528X).

## Acknowledgement

We thank Tommaso Mari and Marie-Luise Kirchner for scientific discussions and JPT peptide technologies for providing peptide membranes at cost. E.R. was supported by a fellowship of the Berlin School of Integrated Oncology (BSIO).

## Supplemental Figures Legends

**Supplemental Figure 1:**
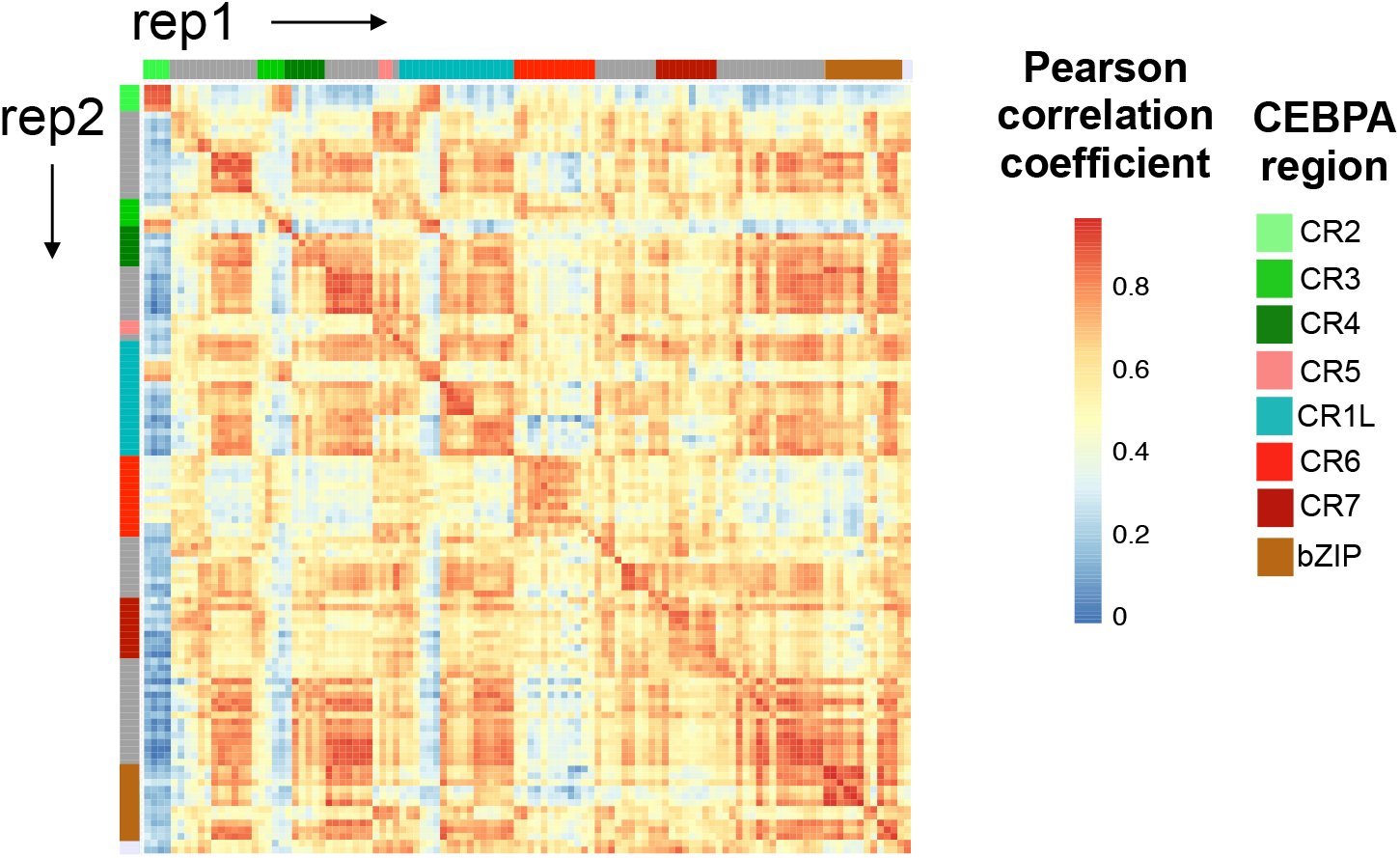
Pearson correlation matrix of PRISMA replicates. Pearson correlation of log2 LFQ values in each PRISMA spot (114 spots). Replicate 1 is plotted against replicate 2. Annotation bars indicates conserved regions of CEBPA.

**Supplemental Figure 2:**
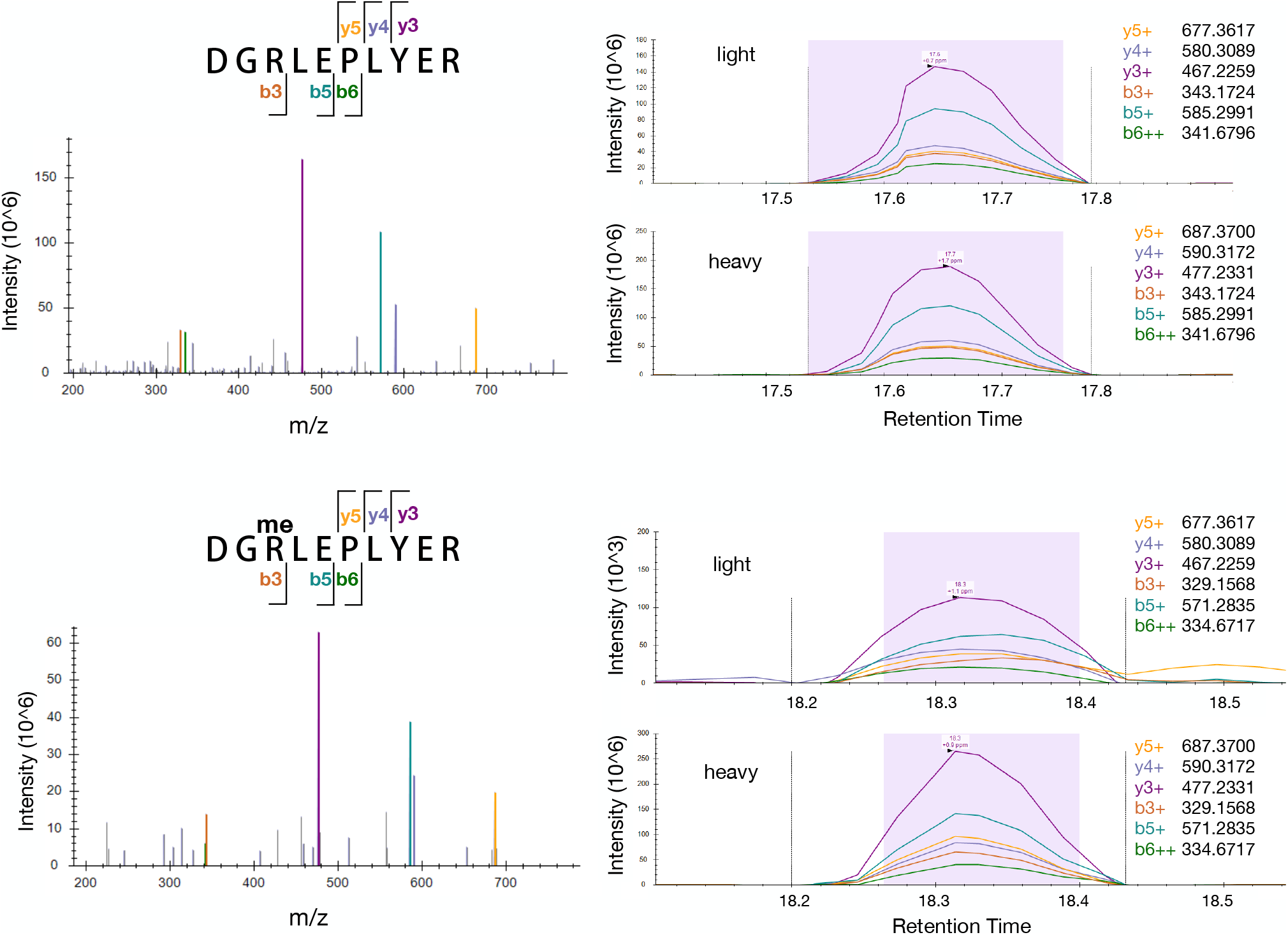
Parallel reaction monitoring detects CEBPA methylation at R142. Parallel reaction monitoring (PRM) of unmodified (top) and methylated (bottom) peptides spanning R142. A heavy peptide standard isotopically labeled at the C-terminal arginine was used to confirm the identity of the peptide. Left: MS2 spectrum of heavy peptide standard; right: elution profile of MS2 fragments of light and heavy peptides. PRM data was analyzed with Skyline.

**Supplemental Figure 3:**
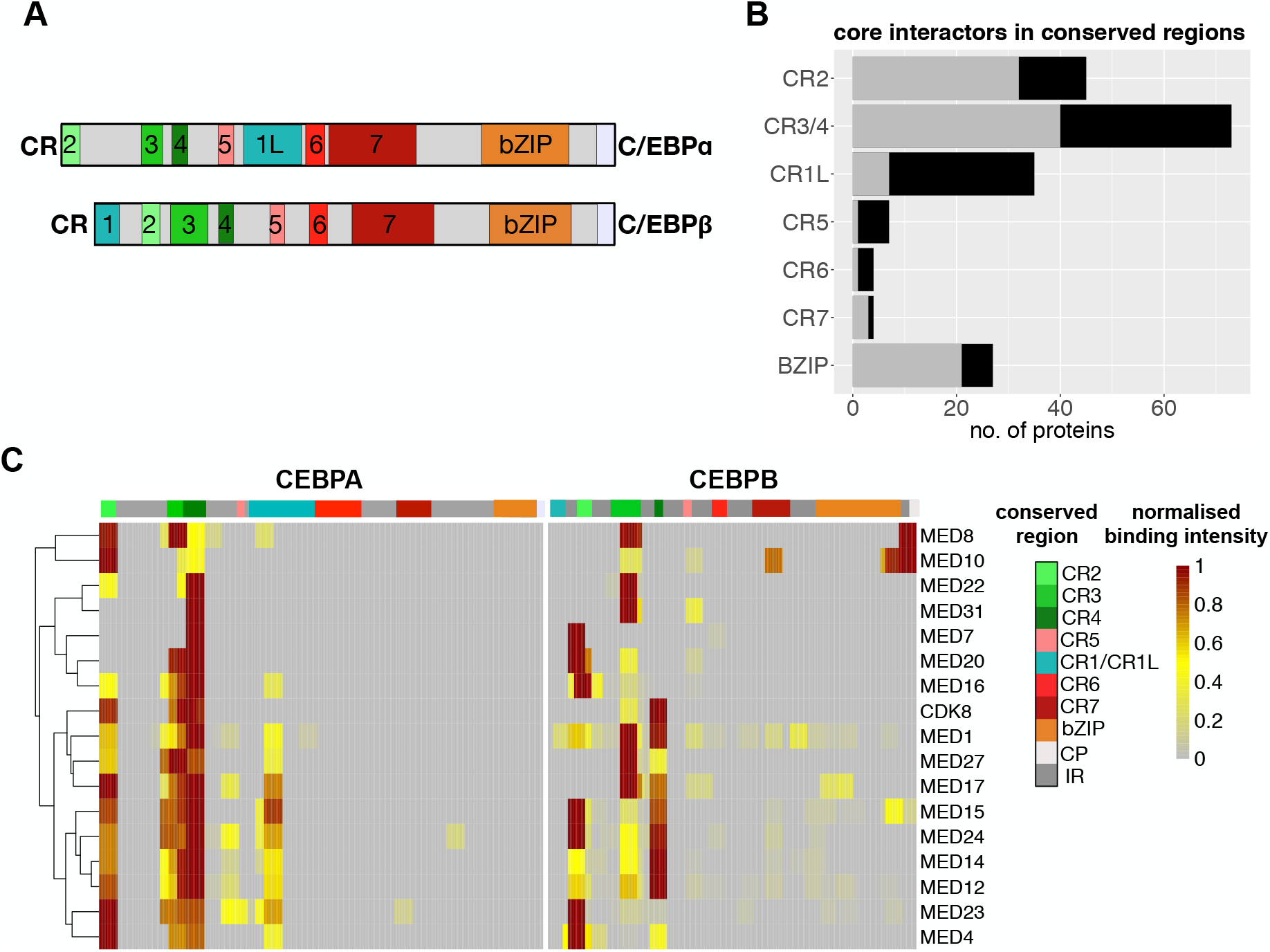
CEBPA and CEBPB share interactors in homologous regions. A: Shared homology of conserved regions in CEBPA and CEBPB. B: Number of high-confidence CEBPA interactors per conserved region in CEBPA as detected by PRISMA and BioID (black bars). Grey bars represent interactors that were also identified as interactors in homologues regions in CEBPB (Dittmar et al., 2019). C: Extracted PRISMA binding profiles of Mediator complex subunits binding to CEBPA and CEBPB (right). Annotation bar on top indicates conserved regions.

**Supplemental Figure 4:**
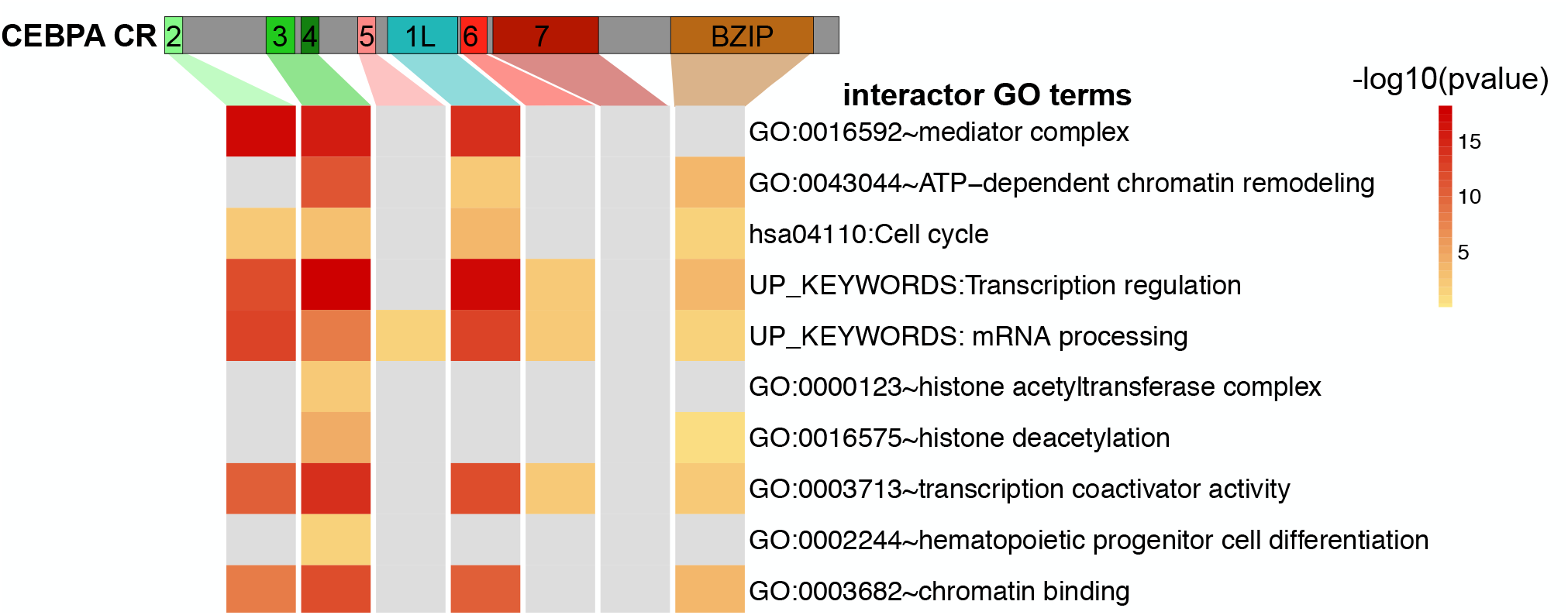
GO term enrichment of mapped CEBPA interactors. PRISMA CEBPA interactors confirmed by BioID were subjected to GO term analysis using the DAVID tool. Informative significant GO terms (p value < 0.05) are displayed. Grey indicates no significant enrichment. No GO terms were significantly enriched in CR7 binders.

**Supplemental Figure 5:**
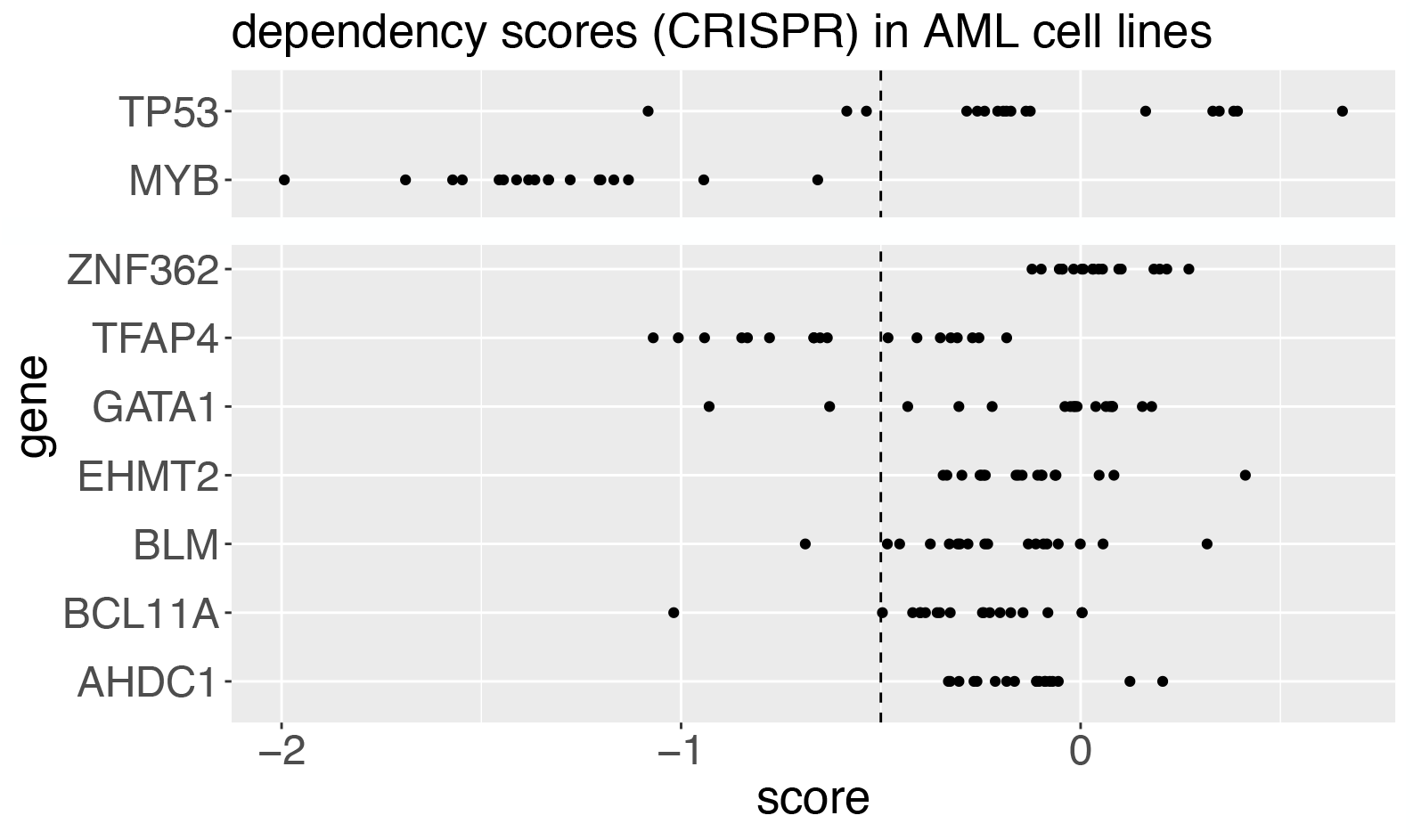
CEBPA P30 specific interactors may represent therapeutic targets in AML. Dependency scores from CRISPR knockout screens in AML cell lines extracted from the DepMap portal. Scores of P30 specific interactors are displayed. Known tumor-suppressor P53 and oncogene MYB are plotted on top as a reference.

